# Phylogeny of the stink bug tribe Chlorocorini (Heteroptera, Pentatomidae) based on DNA and morphological data: the evolution of key phenotypic traits

**DOI:** 10.1101/2020.02.20.957811

**Authors:** Bruno C. Genevcius, Caroline Greve, Samantha Koehler, Rebecca B. Simmons, David A. Rider, Jocelia Grazia, Cristiano F. Schwertner

## Abstract

Pentatomidae is the third largest family of true bugs, comprising over 40 tribes. Few tribes have been studied in a phylogenetic context, and none of them have been examined using molecular data. Moreover, little is known about the evolution of key morphological characters widely used in taxonomic and phylogenetic studies at multiple levels. Here, we conduct a phylogenetic study of the tribe Chlorocorini (Pentatominae) combining 69 morphological characters and five DNA loci. We use the inferred phylogeny to reconstruct the evolution of key morphological characters such as the spined humeral angles of the pronotum, a dorsal projection on the apices of the femora and characters of male genitalia. We provide solid evidence that the tribe as currently recognized is not monophyletic based both on DNA and morphological data. The genera *Arvelius* Spinola and *Eludocoris* Thomas were consistently placed outside of the Chlorocorini, while the remaining genera were found to form a monophyletic group. We also show that nearly all morphological diagnostic characters for the tribe are homoplastic. The only exception is the development of the hypandrium, which, contrary to expectations for genital traits, showed the slowest evolutionary rates. In contrast, the most rapidly evolving trait is the length of the ostiolar ruga, which may be attributed to selection favoring anti-predatory behavior and other functions of its associated scent glands. Lastly, we also provide a preliminary glimpse of the main phylogenetic relationships within the Pentatomidae, which indicates that most of the included subfamilies and tribes are not monophyletic. Our results suggest that the current subfamily-level classification of Pentatomidae is not adequate to reflect its evolutionary history, and we urge for a more complete phylogeny of the family.

## INTRODUCTION

Pentatomidae are the third largest family of true bugs (Hemiptera, Heteroptera). With nearly 5,000 species and over 900 genera, pentatomids are distributed in all terrestrial biomes, except Antarctica (Grazia *et al*., 2015). Numerous species are regarded as major pests of several crops around the world, being responsible for losses of millions of dollars each year (McPherson, 2018). They exhibit a plethora of anatomical and behavioral characteristics which make the group an interesting model for evolutionary and ecological questions. Examples of these features include a variety of feeding habits (Weirauch *et al*., 2018), aposematism (Paleari, 2013), exaggerated sexual traits (McLain, 1981) and parental care (Requena *et al*., 2010). However, evolutionary studies addressing these topics are practically unfeasible with pentatomids due to the absence of phylogenetic hypotheses for major groups. While the position of the Pentatomidae was secondarily explored in studies focusing on other pentatomoids (e.g. Wu et al. 2016; Liu et al. 2019), our knowledge about the lineages that compose the family and the relationships among them remain elusive. Even less is known about the evolution of anatomical and behavioral characteristics mentioned above.

The Pentatomidae are currently divided into nine or ten subfamilies, depending on the classification hypothesis (Schuh & Slater, 1995; Grazia *et al*., 2015; Rider *et al*., 2018). Most subfamilies are arguably monophyletic as they exhibit sets of unique and remarkable anatomical features not present in any other group (Rider, 2000). For example, the Asopinae display head and mouthpart modifications that most likely represent a single-origin adaptation to predatory lifestyles (Parveen *et al*., 2015). The exception is the most diverse subfamily, Pentatominae, the monophyly of which has been broadly questioned (Grazia *et al*., 2008a, 2015). The classification within this group has been called “chaotic” (Rider, 2000) and currently comprises over 40 tribes that encompasses all genera that do not fit in the other subfamilies. Few tribes of Pentatomidae have been studied in a phylogenetic context (e.g. Campos and Grazia 2006; Bernardes et al. 2009; Schwertner and Grazia 2012), and none of these studies used molecular data. The current classification of pentatomid tribes and subfamilies is still based on traditional taxonomic studies in the absence of a phylogenetic context. The reliability of the taxonomic characters for identifying natural groups has not been assessed with independent datasets. The availability of molecular data for species of Pentatomidae has increased during recent years; however, taxon sampling is extremely biased towards Asian and European species (e.g. Yuan et al. 2015; Wu et al. 2016; Liu et al. 2019). Establishing phylogenetic relationships for the pentatomid fauna of the New World is considered paramount for developing a more accurate classification for the family, and for better understanding the evolution of stink bugs.

Out of the tribes of Pentatomidae that occur exclusively in the New World, the Chlorocorini stand out as the most speciose (Rider *et al*., 2018). The tribe is comprised of 78 species organized into eight genera: *Arvelius* Spinola (18 spp.), *Chlorocoris* Spinola (24 spp.), *Chloropepla* Stål (13 spp.), *Eludocoris* Thomas (1 sp.), *Fecelia* Stål (4 spp.), *Loxa* Amyot and Serville (10 spp.), *Mayrinia* Horváth (4 spp.), and *Rhyncholepta* Bergroth (4 spp.). *Chlorocoris* is the most diverse genus, and it is the only one divided into sub-genera: *Chlorocoris (Arawacoris), Chlorocoris (Chlorocoris)* and *Chlorocoris* (*Monochrocerus*). Several authors previously suggested close relationships among some of the genera, formerly placing them in the tribe Pentatomini (Becker & Grazia, 1971; Rolston & McDonald, 1984). Stål (1868) described *Chloropepla* and keyed the genus together with *Chlorocoris* and *Loxa*; later, he also included *Fecelia* in his key (Stål, 1872). More recently, Grazia (1968, 1976) suggested *Chlorocoris, Chloropepla*, *Loxa*, *Mayrinia* and *Fecelia* to be related based on the general morphology (e.g. body coloration, head and general body shape), highlighting the presence of spined humeral angles and a dorsal apical projection on each femur as the main diagnostic characters) (Fig. 1); however, femoral projections are not found in *Chlorocoris* (Eger, 1978; Thomas, 1985). The genera *Arvelius, Eludocoris* and *Rhyncholepta* were later added to the group based on the presence of at least some of morphological features described as characteristic of the tribe, such as the humeral angles projected and the tapering apices of the juga (Becker & Grazia, 1971; Thomas, 1992; Greve *et al*., 2013; Kment *et al*., 2018; Rider *et al*., 2018). However, these eight genera were only recently recognized formally as a distinct tribe within the Pentatominae (Rider *et al*., 2018).

**Figure 1.**
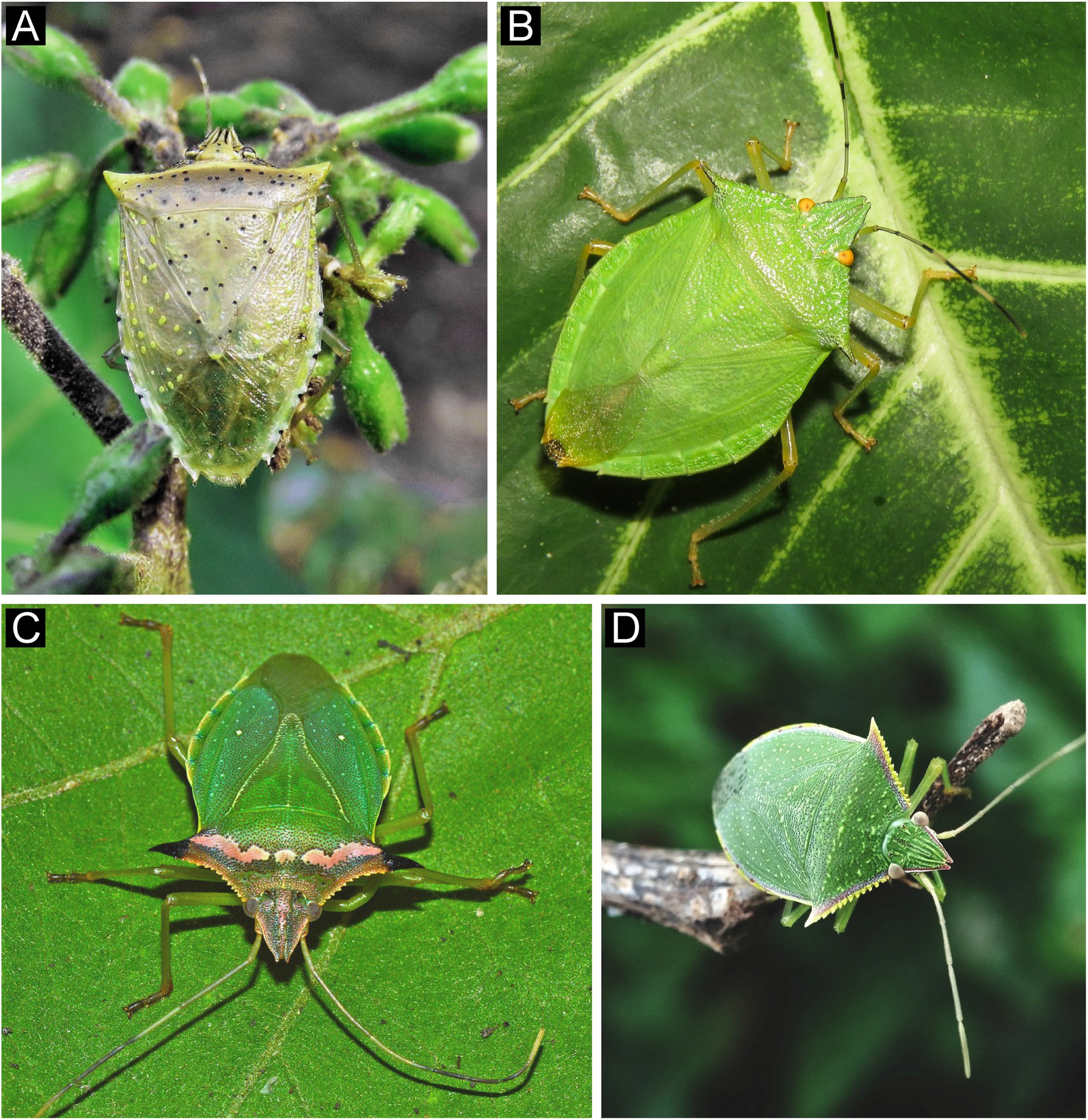
Examples of morphological diversity within the Chlorocorini. (A) *Arvelius albopunctatus* (DeGeer) (photo by Roger Rios Dias), (B) *Chlorocoris complanatus* (Guérin-Méneville) (photo by Diogo Luiz), (C) *Fecelia nigridens* (Walker) (photo by Francisco Alba Suriel) and *Loxa* sp. (photo by Maurino André).

The tribe has been considered to be monophyletic based on several characteristics found on nearly all body parts (Greve *et al*., 2013; Rider *et al*., 2018). Some of the defining characters are the triangular head, the spined humeral angles on the pronotum, the dorsal projections on the apices of the femora, the absence of an abdominal spine, and the presence of a well-developed pair of projections (“hypandrium”) in the male genital capsule. There are also additional features that, in combination, are diagnostic for species of the Chlorocorini: a depressed body, anterior pronotal margins with conspicuous denticles, short ostiolar rugae, and a medially-carinate mesosternum (Becker & Grazia, 1971; Greve *et al*., 2013; Rider *et al*., 2018). Nevertheless, it is noteworthy that many of these characteristics are either absent in part of the Chlorocorini species (e.g. the aforementioned apical projection of the femur) or exhibited by other groups outside of the tribe (Barcellos & Grazia, 2003; Bernardes *et al*., 2009). Interestingly, some of these structures are broadly used in taxonomic and phylogenetic studies of Heteroptera and Hemiptera as a whole. Nevertheless, little is known about their patterns and processes of diversification in Pentatomidae, making it unclear if these structures are indeed informative for the evolutionary history of these bugs. Only few studies have attempted to reconstruct the evolution of such key morphological characters in evolutionary time (Genevcius *et al*., 2017). As a result, we still have a superficial comprehension of the tempo and mode of morphological diversification in these bugs.

Herein, we conducted a phylogenetic study of the tribe Chlorocorini including representatives of all genera and the three subgenera of *Chlorocoris* within the ingroup taxa. This is the first phylogenetic analysis within the Pentatomidae at the tribal level using a combination of morphological and molecular data. By integrating partial sequences of five molecular markers and 69 morphological characters, we first aimed to test the monophyly of the tribe, recognize lineages within the Chlorocorini and determine its phylogenetic position within the family. Second, we reconstructed the evolution of key morphological characters to determine their patterns of diversification within the tree and determine their levels of homoplasy. Additionally, we investigated the tempo of diversification of these characters by estimating their evolutionary rates in a model-based approach. We discuss broader implications of our findings for studies on morphological evolution as a whole and for the classification of Pentatomidae.

## MATERIALS AND METHODS

### Taxon sampling

Our analyses included 37 terminal taxa. The ingroup sampling was comprised of twelve representatives of the eight genera of Chlorocorini and the three subgenera of *Chlorocoris*. As outgroups, we included 23 species from four subfamilies of Pentatomidae that occur in the Neotropics (Asopinae, Discocephalinae, Edessinae and Pentatominae) and two species of closely related families (Scutelleridae and Acanthosomatidae). Our outgroup choices are carefully designed to include genera that also exhibit some of the characters that have been considered to be synapomorphic within the Chlorocorini, with emphasis placed on the New World fauna.

### DNA markers and data acquisition

Genomic DNA was extracted from thoracic muscle of adults previously stored in 100% ethanol and preserved at −80 °C. We used the DNeasy Blood & Tissue kit (QIAGEN, Valencia, CA), following the manufacturer’s protocol. Partial sequences of five loci were amplified, comprising four ribosomal gene regions (16S rDNA, 18S rDNA, 28S D1 rDNA, 28S D3-D5 rDNA) and one mitochondrial protein-coding gene (COI). Primer sequences used in amplification and sequencing are listed in table S1. Polymerase chain reaction (PCR) was performed using the Platinum^®^ Taq DNA Polymerase kit (Invitrogen, Brazil) considering a final volume of 25 μl and 1× PCR buffer, 10 mM dNTP, 2.5 μl of 3.0 mM MgCl2, 0.4 μM of each primer, and 20 – 50 ng of template DNA. PCR programs were the same for all markers (except for the annealing temperature): 94°C for 1 min, 35 cycles of 94°C 30 s, 45–55°C 30 s (Table S1), 72°C 1 min, and 72°C for 10 min. Amplification results were visualized using gel electrophoresis (1% gel agarose) with the SyberSafe gel staining and UV illuminator. Purified DNA was sequenced by Macrogen, Inc (Seoul, South Korea). Sequences were deposited in NCBI-GenBank (https://www.ncbi.nlm.nih.gov/genbank/) and access codes are provided in the supplementary material (Table S2).

### Sequence alignment and phylogenetic analyses

Sequences were assembled and edited in GENEIOUS 7.1.7 (Biomatters, Auckland, New Zealand). Alignments were conducted individually in the online v. 7 of MAFFT using the G-INS-i algorithm and default parameters, and were checked manually for inconsistencies. Sequence-matrix 1.8 was used to combine individually aligned molecular markers into a single partitioned dataset.

Molecular and morphological datasets were analyzed, both separately and in combination in a total evidence partitioned analysis. Bayesian Inference was our main criterion for phylogenetic analyses, but we also ran a Maximum Likelihood analysis on the combined dataset to check for consistency. Different evolutionary models were allowed for each DNA marker partition, while the Mk model was constrained for the morphological partition. Model selection for DNA markers was run using ModelFinder (Kalyaanamoorthy *et al*., 2017) through the IQ-TREE software v.1.6.9 (Nguyen *et al*., 2015), using the BIC criterion and restricting to models available in Mr. Bayes (Ronquist *et al*., 2012). Bayesian phylogenetic analyses were run in Mr. Bayes v.3.2.6 (Ronquist *et al*., 2012) through the CIPRES web portal (http://www.phylo.org) with the following parameters: 30 million generations, two independent runs, four chains, default priors, and sampling trees every 3000 generations. Convergence among runs were checked by confirming that the average standard deviation of split frequencies reached < 0.01, and we ensured that effective sample sizes (ESS) were at least 200. Posterior distributions of parameter estimates were visualized in TRACER v.1.6.0. We discarded the first 10% of the generations as burn-in and constructed a 50% majority rule consensus from the remaining trees. Maximum Likelihood analyses were run in IQ-TREE using the same transition models as in Mr. Bayes and 1000 replicates of ultra-fast bootstrap. Phylogenetic trees were visualized and edited in FigTree v.1.4.0 (http://tree.bio.ed.ac.uk/software/figtree).

### Morphological characters

Morphological characters were either reinterpreted from the literature or described for the first time, totaling 69 morphological features of the entire body: head, thorax, abdomen and the genitalia (character description in Table S3). Terminology follows Greve et al. (2013) and references therein; characters based on literature are listed and indicated (Table S3). Newly described characters were analyzed using a stereo microscope Leica M205C. Coding of morphological characters can be found in Table S4.

### Evolution of key morphological characters

We recognized *a priori* eight morphological characters that have been proposed as key characters for the recognition of species of Chlorocorini (Rider *et al*., 2018). The same structures are consistently present in phylogenetic and taxonomic studies of Pentatomidae and related groups. To analyze the diversification patterns of these characters within Chlorocorini, we conducted a maximum likelihood ancestral state reconstruction using MESQUITE v. 3.0.4. We report the likelihood of the ancestral states at selected branches where evolutionary changes are more likely to have happened. These analyses allowed us to determine the ancestral and derived conditions and to evaluate whether diagnostic characters are phylogenetically informative.

We employed Bayesian trait modeling to estimate the evolutionary rate of change for the selected characters in BayesTraits v3.0 (Pagel & Meade, 2006). We used a reversible-jump Markov-Chain Monte-Carlo (rjMCMC), which provides posterior probabilities of evolutionary rates of categorical traits not restricted to a single evolutionary model (Pagel & Meade, 2006). Markov-chains were run for 10 million iterations, sampling every 100 thousand iterations after 1 million samples of burn-in. The prior was an exponential distribution with the mean seeded from a uniform distribution ranging between 0 and 2. For each trait, we concatenated the posterior probabilities of each possible transition and reported these distributions.

## RESULTS

Bayesian analysis of the morphological data alone resulted in a consensus tree with low posterior probabilities overall and several polytomies, especially at deeper nodes (Fig. S1). The tribe Chlorocorini did not emerge as monophyletic because *Arvelius* was placed outside the tribe, although the position of the tribe was largely uncertain and the support of *Arvelius* + *Taurocerus* is moderate (Fig. S1). The remaining Chlorocorini formed a monophyletic group with poor support. The only major groups recognized with high support in the morphological tree were the Discocephalinae (p.p. = 1), the Edessinae (p.p. = 1), the Asopinae (p.p. = 1), and a group of Neotropical genera currently included in the tribe Carpocorini (p.p. = 0.95).

Evolutionary models for each molecular marker selected in ModelFinder are provided in Table S5. In summary, the simple two-parameter Kimura model was selected for the nuclear regions (18S and 28S), while more complex models were selected for the mitochondrial regions (GTR+I+G for COI and HKY+I+G for 16S). Analysis of the combined molecular markers showed substantially improved posterior probabilities and overall better resolution than the morphological tree (Fig. S2). The monophyly of Chlorocorini had even less support in this analysis (Fig. S2) as *Arvelius*, *Chlorocoris* and *Eludocoris* were phylogenetically associated with other tribes or subfamilies. The position of *Arvelius* was congruent with the morphological analysis, although support levels ranged from moderate to high. The genus was placed outside of the Chlorocorini and as closely related to *Taurocerus edessoides* (Spinola). However, the position of *Chlorocoris*, which was consistently monophyletic, and *Eludocoris grandis* were incongruent between the morphological and molecular topologies, appearing in the latter as closely related to the Edessinae and *Ocirrhoe*, respectively. The remaining genera currently placed in the Chlorocorini formed a monophyletic group with low support as in the morphological analysis, although relationships within this group were significantly different from the morphological analysis (Figs. S1-S2). In contrast with the morphological analysis, only the Neotropical Carpocorini and the Nezarini were recovered as monophyletic (p.p. = 1 and 0.63, respectively). Neither Edessinae or Discocephalinae did emerge as monophyletic, while the subfamily Asopinae (monophyletic, p.p. = 1) was phylogenetically related to species from tribes of Pentatominae (p.p. = 0.91; Fig. S2).

Although morphological and molecular datasets were only partially congruent, the total evidence analyses had highest resolution and greater node support (Fig. 2). Bayesian (BI – Fig. 2) and Maximum Likelihood (ML – Fig. S3) trees were highly similar, supporting the same major groups and, most importantly, indicating the tribe Chlorocorini as polyphyletic. *Arvelius* was recovered as more closely related to *Taurocerus* (p.p. = 1) and other lineages of Pentatominae + Asopinae (p.p. = 1). Both analyses congruently indicate that *Eludocoris* was phylogenetically associated with the Edessinae, *Ocirrhoe* and *Myota*, as well as *Rhyncholepta* as the sister genus of the remaining Chlorocorini. Because BI and ML trees were largely congruent and our main conclusions are the same, we report detailed results only for the Bayesian analysis below.

**Figure 2.**
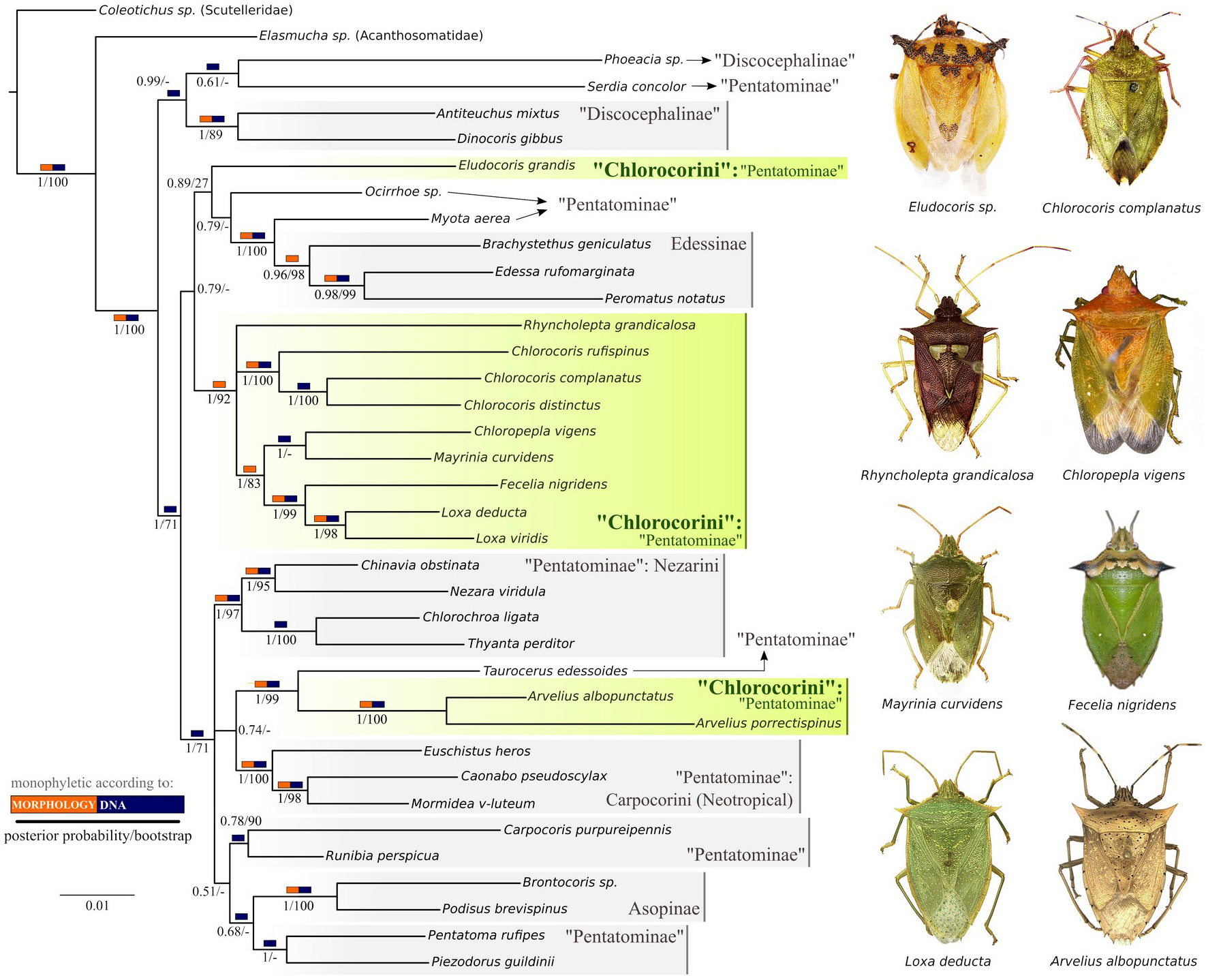
Bayesian majority consensus tree constructed using combined morphological (69 characters) and DNA sequence data (16S rDNA, 18S rDNA, 28S D1 rDNA, 28S D3-D5 rDNA and COI mitDNA), along with the dorsal habitus of all genera of Chlorocorini. Rectangles above branches indicate whether the clade is supported by morphological and/or molecular data, while numbers below branches are posterior probabilities and bootstrap supports from the maximum likelihood analysis.

The total evidence Bayesian tree displays relationships found in both morphological and molecular topologies, with more similarities with the molecular tree. Only three clades in the total evidence tree are supported only by morphological evidence (orange rectangles in Fig. 2: Edessinae, *Rhyncholepta* + the remaining Chlorocorini (excluding *Arvelius* and *Eludocoris*) and *Chloropepla + Mayrinia + Fecelia* + *Loxa*. In contrast, ten clades are supported exclusively by DNA data (e.g. the relationships between *Chloropepla* and *Mayrinia* and between *Chlorocoris complanatus* and *Chlorocoris distinctus*) (Fig. 2). Relationships at deeper nodes were also improved in the total-evidence tree. For example, the Discocephalinae and *Serdia* Stål (currently placed in the Pentatomini) are strongly supported as the sister lineage of the remaining pentatomids (Fig. 2).

The polyphyly of Chlorocorini was also corroborated by the total-evidence analysis. The tribe emerged, again, as polyphyletic due to the position of *Arvelius* and *Eludocoris* (Fig. 2). *Arvelius* was closely related to *Taurocerus* and other tribes of Pentatominae, while *Eludocoris* was the sister group of Edessinae + *Myota* + *Ocirrhoe*. The remaining genera formed a well-supported monophyletic group, which could be split into three groups: the three subgenera of *Chlorocoris*, the genus *Rhyncholepta*, and the clade *Loxa* + *Fecelia* + *Mayrinia* + *Chloropepla*. The sister group of Chlorocorini (excluding *Arvelius* and *Eludocoris*), which was ambiguous in the separated analyses of each type of data (morphological versus molecular), was the Edessinae plus *Myota* Spinola and *Eludocoris*. This relationship had moderate support.

Reconstructed ancestral states of eight selected characters (Fig. 3) revealed that all genera of the Chlorocorini (excluding *Arvelius* and *Eludocoris*) share several synapomorphies (characters 3^1^, 5^1^, 22^1^, 26^0^, 34^1^, 57^1^ and 65^1^). These synapomorphies include characters of the head, thorax, abdomen and genitalia (Fig. 3), such as a long ostiolar ruga (character 26^0^), dorsal apical projections in the femora (34^1^) and a well-developed conjunctiva (65^1^). The only character proposed as diagnostic that was not synapomorphic for the tribe was the shape of juga (7^0^). Nevertheless, our results also show that most of the key characters have changed multiple times across the phylogeny.

**Figure 3.**
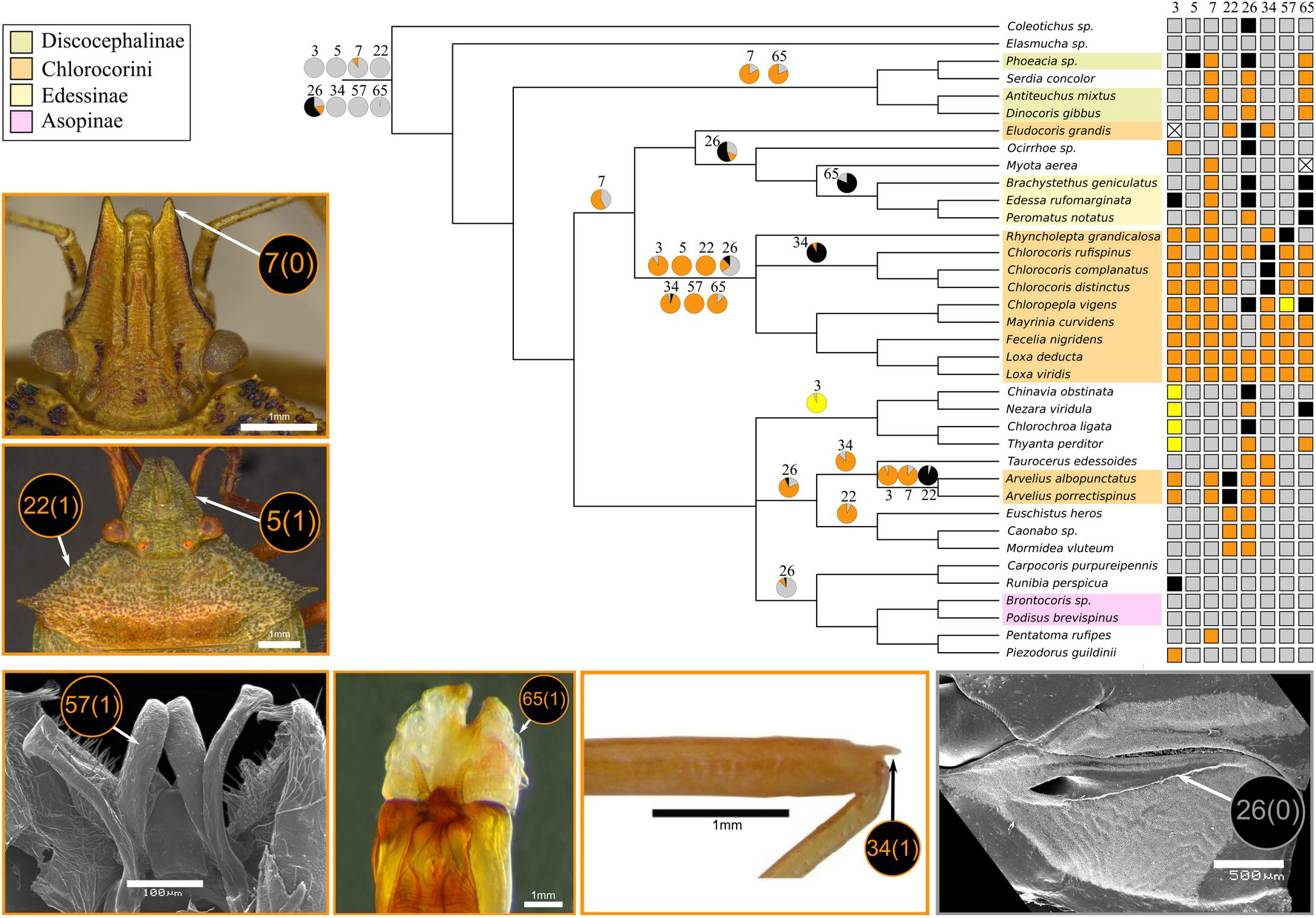
Maximum likelihood ancestral states of the key morphological characters of Chlorocorini reconstructed over the total evidence bayesian tree (converted to cladogram for visualization). Likelihood of ancestral states are shown as pie charts only in the nodes where evolutionary changes are likely to have happened (changes in terminal branches are omitted). Scoring for each taxon are exhibited as colored rectangles, where crossed rectangles are inapplicable states (gray = state 0, yellow = 1, orange = 2, black = 3, crosses = missing/not applicable).

The analyses of evolutionary rates indicate that the length of the ostiolar ruga is to most labile character, showing a five-fold increase in the rate of evolution in comparison to the rate of change displayed by the margin of the head (Fig. 4). The second structure with faster evolutionary rates was the length of the juga. In contrast, the development of the hypandrium was the slowest evolving character, with a mean rate of 0.78. All other characters showed similar intermediate values, ranging between 1.36 and 2.32 (Fig. 4).

**Figure 4.**
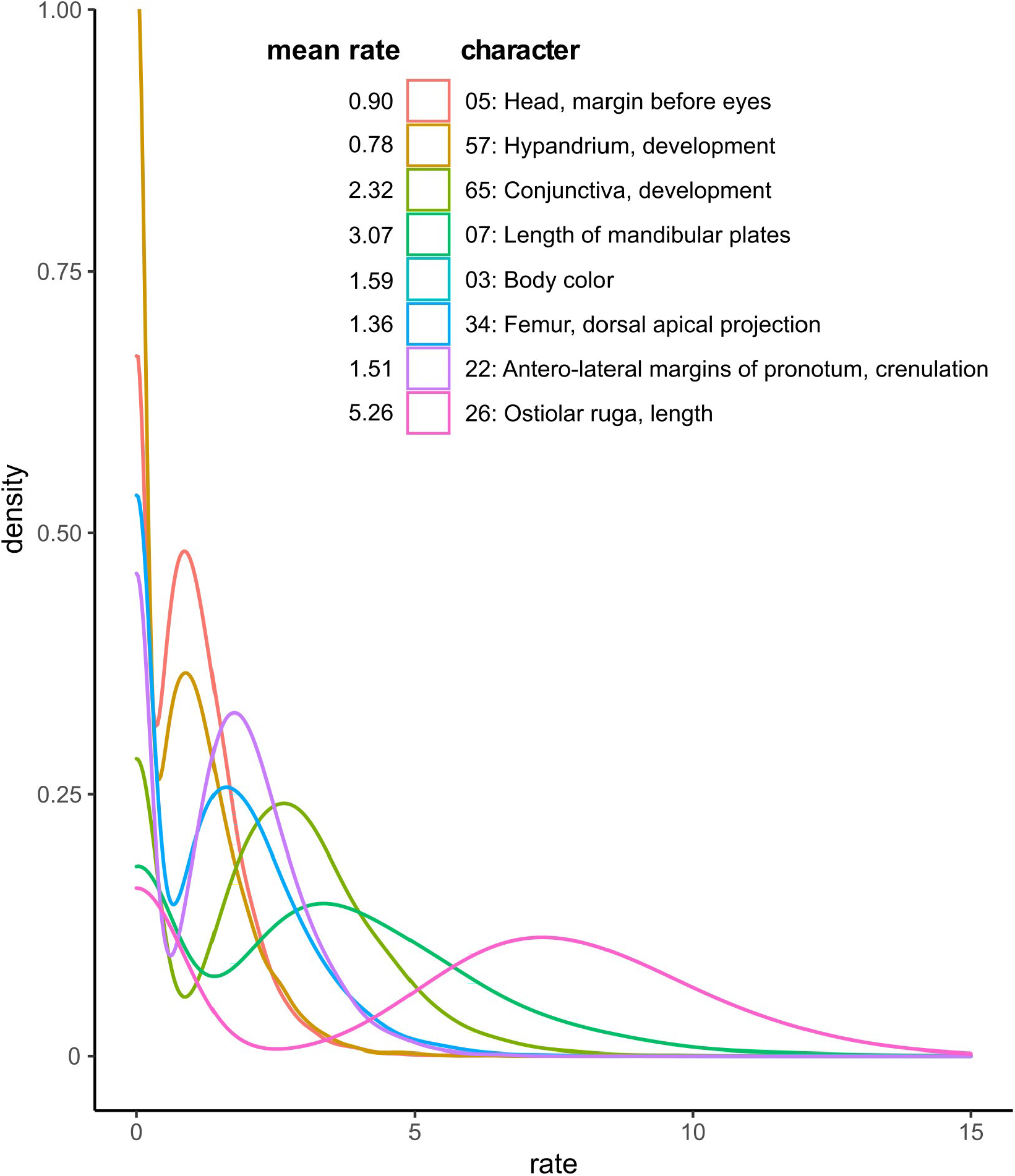
Posterior distributions (kernel density plots) of estimated evolutionary rates for the eight key morphological characters.

## DISCUSSION

### Monophyly of Chlorocorini

Our study proposes the first phylogenetic hypothesis for subfamilies and tribes of Pentatomidae using both molecular and morphological data. The morphological data analyzed alone provided good resolution for families and tribes, but also left the backbone of the tree mostly unresolved (Fig. S1). In contrast, the molecular tree has much better resolution from tips to the root, even though many clades showed modest to poor posterior probabilities (Fig. S2). When combined, morphological and molecular data resulted in the tree with the best resolution and higher mean support levels.

Our analyses refuted the hypothesis of monophyly for the Chlorocorini as currently recognized by Rider *et al*. (2018). The polyphyly of the tribe is owing to the position of two genera: *Arvelius* (in all analyses) and *Eludocoris* (except in the morphological analysis). It should be noted that (Rider *et al*., 2018) indicated that *Arvelius* might not belong to this tribe, based on morphological characters, especially the armed nature of the abdominal base. The genus consistently emerged in our results as the sister group of *Taurocerus* Amyot & Serville; both were more closely related to other genera of Pentatominae than to genera of Chlorocorini. This result is not altogether surprising as both Brailovsky (1981) and Grazia & Barcellos (2005) highlighted similarities between the two genera, including the apical projection on each femur, the abdominal spine present and opposed to the elevated metasternum, and the morphological aspect of the phallus. According to Thomas (1992), *Eludocoris* should be placed near *Loxa* and allied genera (*Chlorocoris*, *Chloropepla* and *Fecelia*) based primarily on the large body size, the elongate and depressed body shape, the greenish coloration when alive, and the presence of an apical projection on each femur. This hypothesis agrees with our morphological topology. On the other hand, the molecular data indicate that *Eludocoris* may be closely related to the Australian genus *Ocirrhoe* (which is currently placed in the tribe Rhynchocorini), and both genera were the sister group of the remaining Pentatominae. We caution, however, that these possible placements are tentative at best, and further studies are needed to verify these observations.

### Position of Chlorocorini and relationships among genera

While the non-monophyly of Chlorocorini as currently recognized was strongly supported, the position of its genera among the other lineages of the Pentatomidae was less consistent. The morphological tree exhibited a large basal polytomy that did not provide certainty for identifying the sister group of Chlorocorini excluding *Arvelius* and *Eludocoris*. In the molecular analysis, the genus *Chlorocoris* was closely related to the Edessinae, while the remaining genera (except *Arvelius*) were the sister group of the Pentatominae (including Asopinae). Lastly, results of the combined analysis were partially congruent with the molecular analysis. Accordingly, all genera of Chlorocorini (except *Arvelius*) were grouped with the Edessinae, and this clade was the sister lineage of a clade including the Asopinae and the remaining Pentatominae.

Although the position of Chlorocorini was poorly supported with either DNA or morphological data analyzed individually, there were some consistent relationships between the two datasets. None of the genera (except *Arvelius*) were placed within the Pentatominae, which provides no certainty for including the Chlorocorini as a tribe of Pentatominae as in the current classification of the family (Greve *et al*., 2013; Rider *et al*., 2018). We conclude that the genera *Chlorocoris, Chloropepla, Fecelia, Loxa, Mayrinia* and *Rhyncholepta* indeed compose a distinct evolutionary lineage. This group should either comprise a new subfamily or be incorporated with *Eludocoris* into the subfamily Edessinae. We do not seek to formally propose one of these changes here because of the poor resolution of these clades and because our taxon sampling is limited (especially with respect to the European, Asiatic and Australian faunas). Furthermore, there is not a single morphological synapomorphy for the clade comprising the Edessinae and the Chlorocorini, which would require closer inspection of the morphology focusing on an alternative level of analysis. Therefore, to allow for such taxonomic decisions, we urge for a complete phylogeny of the Pentatomidae with datasets including more morphological and molecular data.

Overall, relationships within Chlorocorini were more stable than the tribe’s position within the subfamily. Furthermore, some relationships among genera are in close agreement with the literature, suggesting that traditional taxonomic characters are more reliable at the generic level. All analyses supported the monophyly of the genera *Arvelius, Chlorocoris* and *Loxa*, as well as the monophyletic group *Fecelia + Loxa*. The latter was also consistently associated with *Chloropepla* and *Mayrinia*, although *Rhyncholepta* may also fall within this group according to DNA data (Fig. S2). The association among *Chloropepla*, *Fecelia*, *Loxa* and *Mayrinia* was not surprising as it has been repeatedly suggested in the literature based on shared features of the head, pronotum and scutellum shape (Grazia, 1972; Rolston & McDonald, 1984; Greve *et al*., 2013). The ambiguous position of *Rhyncholepta* may be explained by its highly autapomorphic features. The genus possesses several unique characteristics in the tribe, such as the body color (after death), longer antennae, larger eyes, presence of an abdominal tubercle, and several genital features (Becker & Grazia, 1971; Kment *et al*., 2018). Some of these characteristics have been included in our analyses (e.g. characters 35, 57), which may have supported its position as sister of the remaining species within Chlorocorini.

### Evolution of key morphological characters

Our outgroup selection provided a robust sample that is representative of the Neotropical lineages which also exhibit the diagnostic features of the Chlorocorini. This careful outgroup choice enabled us to confidently investigate both the monophyly of the Chlorocorini and also the amount of homoplasy found in traditional diagnostic features. We show that all characters that group the genera of Chlorocorini in the taxonomic literature (Becker & Grazia, 1971; Grazia *et al*., 2008b; Rider *et al*., 2018) have some level of homoplasy, with the exception of the development of the hypandrium. This character showed the slowest evolutionary rates (Fig. 4), changing only four times across the phylogeny despite having four states (Fig. 3). In principle, this may seem unexpected because genital traits are usually the fastest evolving traits in animals with internal fertilization (Klaczko *et al*., 2015; Genevcius *et al*., 2017). However, one should note that these characters have been delineated and chosen to be informative at higher levels. Therefore, although the male genitalia may indeed evolve fast in Pentatomidae (Genevcius *et al*., 2017), our data indicate that more conserved and phylogenetically informative characters can also be found in this character system.

Based on our analyses, the most homoplastic character was the length of the ostiolar ruga (Fig. 3). The ancestral form most likely was a longer ruga, which experienced length decreases and subsequent reversions multiple times during the evolution of Pentatomidae (Fig. 4). Our results agree with other comparative studies that revealed strong interspecific variability for these characters in other groups of pentatomids (Kment & Vilímová, 2010; Barão *et al*., 2017). Interestingly, the length of the ostiolar ruga is one of the characters widely used for the classification (and diagnostics) within Pentatomidae (Rider *et al*., 2018), and its homoplastic nature has already been suggested in other phylogenetic studies (Barcellos & Grazia, 2003; Memon *et al*., 2011). This plasticity may be suggestive of an adaptive role for the ostiolar ruga. Although their function is yet poorly understood, the ruga are part of the thoracic scent efferent system. As such, changes in their shape and size may have critical ecological outcomes associated with antipredatory behavior and other functions of the scent glands within Pentatomidae (i.e. intraspecific communication).

The remaining characters that we examined showed intermediate levels of homoplasy and evolutionary rates. In fact, relatively high levels of homoplasy for characters describing features such as head shape and development of projections are not altogether surprising, as these have been shown to be homoplastic in many other phylogenetic studies within Pentatomidae (e.g. Genevcius and Schwertner 2014; Weiler et al. 2016; Bianchi et al. 2017). One likely reason for high levels of homoplasy is that these characters are arbitrary categories of structures with continuous variation; therefore, states for these characters tend to overlap (Cohen, 2012). Most of the diagnostic characters for the Chlorocorini are likely artificial constructs, for example, the shape of the juga (character 7), the length of the ostiolar ruga (character 26), and the degree of development of the conjunctiva (character 65). These results call the value of these characters into question, especially as used to propose a classification of the Pentatomidae based on the evolutionary history of the group. Further phylogenetic and taxonomic studies should take into account that these characters are intrinsically continuous, and their use should be considered with caution.

### Implications for the Pentatomidae classification and future directions

Although not the primary focus of this study, our results also provide a preliminary glimpse of potential major relationships within the Pentatomidae. In agreement with previous studies based on morphology (Gapud, 1991; Thomas, 1992) and DNA (Wu *et al*., 2016; Liu *et al*., 2019), our study corroborate the monophyly and the position of the Asopinae within of Pentatominae (Fig. 2). Edessinae was also found to be monophyletic, while Pentatominae was broadly polyphyletic. This is not surprising given that the Pentatominae is a “catch-all” construct that encompasses all genera not placed in the other subfamilies (Rider *et al*., 2018).

The Discocephalinae can be considered monophyletic only with the inclusion of *Serdia concolor* Ruckes, representing the sister lineage of all remaining Pentatomidae. Although these conclusions are based on a small sample size compared with the diversity of Pentatomidae as a whole, our results indicate that the classification of the Pentatomidae at the subfamily and tribal levels will require a thorough reformulation that reflects the family’s evolutionary history. For example, the tentative results obtained in this study indicate that the subfamily Pentatominae will be valid only with the inclusion of the asopines as a tribe and with the exclusion of Chlorocorini and some other genera currently included in other pentatominae tribes (i.e. *Serdia*). The Chlorocorini could be either raised to the subfamily level or it could be merged with the Edessinae. Other tribes included (i.e. Nezarini, Carpocorini, Catacanthini *[Runibia perspicua* (Fabricius)] and Piezodorini [*Piezodorus guildinii* (Westwood)]), would remain with a new concept of the Pentatominae.

In summary, we provide solid evidence that the currently recognized tribe Chlorocorini and two of the subfamilies of Pentatomidae are not monophyletic. Thus, our results indicate that the current classification of the Pentatomidae (sensu Rider et al. 2018) does not accurately reflect the evolutionary history of the group. The most likely reason for that is that traditional diagnostic characters, many of which are based on continuous variation, show considerable levels of homoplasy. Nevertheless, we emphasize that this study was focused on testing the monophyly and the relationships within the tribe Chlorocorini. Thus, our findings regarding the major relationships within the entire family should be considered, at best, preliminary. It appears that the molecular data at times supports previous morphological studies, but often it is not in congruence. In these cases, we should re-analyze the morphological data, looking for convergent, parallel, or misinterpretations of morphological characters. For the next steps regarding the phylogeny and classification of the family, we encourage a thorough and representative revision seeking to find synapomorphies for early diverging lineages, whose phylogenetic placements were poorly supported in our study. Although the inclusion of molecular data improved phylogenetic resolution overall, many clades, especially those that were unresolved in the morphological analysis, still had low support. This suggests that the inclusion of additional nuclear DNA data possessing low divergence rates will be fundamental to improve phylogenetic understanding of these lineages. Additionally, a reliable classification for the Pentatomidae will only be feasible with a robust taxon sampling scheme representing the global diversity of these insects.

### Updated classification of Chlorocorini

Based on our phylogenetic results, we provide a reclassification of the tribe Chlorocorini, now including six genera and 59 species (see below). The genus *Arvelius* is transferred to the tribe Pentatomini (currently the classification of the related genus *Taurocerus*). The genus *Eludocoris* is considered unplaced owing to its inconsistent position among the three analyses (Figs. 2, S1, S2, S3).

### Checklist of the genera and species of Chlorocorini

**Tribe Chlorocorini** Rider, Greve, Schwertner and Grazia, 2018

#### ***Chlorocoris*** Spinola, 1837

Type species: *Chlorocoris tau* Spinola, 1837, by monotypy.

*Chlorocoris (Monochrocerus) biconicus Thomas, 1985 [NIC, CR, PAN, HON]*

*Chlorocoris (Monochrocerus) championi Distant, 1880 [GTM, MEX]*

*Chlorocoris (Chlorocoris) complanatus (Guérin-Méneville, 1831) [BRA, BOL, PAR, ARG, URU]*

*Chlorocoris (Chlorocoris) deplanatus (Herrich-Schäffer, 1842) [BRA]*

*Chlorocoris (Chlorocoris) depressus (Fabricius, 1803) [COL, VEZ, SUR, BRA, TTO, ECU]*

*Chlorocoris (Chlorocoris) distinctus Signoret, 1851 [USA, MEX, BLZ, NIC, HON, GTM, CR, PAN, COL, ECU]*

*Chlorocoris (Chlorocoris) fabulosus Thomas, 1985 [BRA]*

*Chlorocoris (Monochrocerus) flaviviridis Barber, 1914 [USA]*

*Chlorocoris (Monochrocerus) hebetatus Distant, 1890 [USA, MEX]*

*Chlorocoris (Chlorocoris) humeralis Thomas, 1985 [BOL]*

*Chlorocoris (Monochrocerus) irroratus Distant, 1880 [MEX]*

*Chlorocoris (Chlorocoris) isthmus Thomas, 1985 [CR, PAN, COL, ECU]*

*Chlorocoris (Monochrocerus) loxoides Thomas, 1985 [MEX, GTM, NIC]*

*Chlorocoris (Chlorocoris) nigricornis Schmidt, 1907 [PER, ECU]*

*Chlorocoris (Monochrocerus)* rufispinus Dallas, 1851 [MEX, GTM, HON, CR, PAN]

*Chlorocoris (Monochrocerus) rufopictus Walker, 1868 [MEX]*

*Chlorocoris (Chlorocoris) sanguinursus Thomas, 1985 [SUR, PER, BOL]*

*Chlorocoris (Chlorocoris) sororis Thomas, 1985 [COL]*

*Chlorocoris (Monochrocerus) subrugosus Stål, 1872 [USA, MEX]*

*Chlorocoris (Arawacoris) tarsalis Thomas, 1998 [JAM]*

*Chlorocoris (Chlorocoris) tau* Spinola, 1837 [BRA, ARG, URU]

*Chlorocoris (Chlorocoris) tibialis* Thomas, 1985 [BRA]

*Chlorocoris (Chlorocoris) vandoesburgi* Thomas, 1985 [SUR]

*Chlorocoris (Monochrocerus) werneri* Thomas, 1985 [USA, MEX]

#### ***Chloropepla*** Stål, 1868

Type species: *Loxa vigens* Stål, 1860, by original designation.

*Chloropepla aurea* (Pirán, 1963) [BRA, PER, BOL]

*Chloropepla caxiuanensis* Greve, Schwertner & Grazia, 2013 [VEZ, BRA]

*Chloropepla costaricensis* Greve, Schwertner & Grazia, 2013 [CR]

*Chloropepla dollingi* Grazia, 1987 [GUY, BRA]

*Chloropepla lenti* Grazia, 1968 [VEZ, HON]

*Chloropepla luteipennis* (Westwood, 1837) [BRA]

*Chloropepla paveli* Grazia, Schwertner & Greve, 2008 [BRA, BOL]

*Chloropepla pirani* Grazia-Vieira, 1971 [BOL]

*Chloropepla rideri* Greve, Schwertner & Grazia, 2013 [BRA]

*Chloropepla rolstoni Grazia-Viera, 1973 [FG, BRA, BOL]*

*Chloropepla stysi* Grazia, Schwertner & Greve, 2008 [BRA, ECU]

*Chloropepla tucuruiensis* Grazia & Teradaira, 1980 [BRA]

*Chloropepla vigens* (Stål, 1860) [BRA, ARG, URU]

#### ***Fecelia*** Stål, 1872

Type species: *Loxa minor* Vollenhoven, 1868, by monotypy.

*Fecelia biorbis Eger, 1980 [HAI, DRE]*

*Fecelia minor* (Vollenhoven, 1868) [PUR]

*Fecelia nigridens* (Walker, 1867) [HAI, DRE, TTO]

*Fecelia proxima* Grazia, 1980 [DRE, TTO]

#### ***Loxa*** Amyot & Serville, 1843

Type species: *Cimex flavicollis* Drury, 1773, by subsequent designation (Kirkaldy, 1903)

*Loxa deducta (Walker, 1867) [PAN, VEZ, BRA, CHI, BOL, PAR, ARG, URU]*

*Loxa flavicollis* (Drury, 1773) [USA, MEX, CUB, JAM, CR, HON, BRA, ECU, FG, TTO]

*Loxa melanita Eger, 1978 [GUY, BRA, PER]*

*Loxa nesiotes Horváth, 1925 [CUR, GRE, SLU, PAN, COL, VEZ, GUY, DRE, HAI, SVG]*

*Loxa pallida Van Duzee, 1907 [CUB, DRE, BAH, PUR, JAM, DOM]*

*Loxa parapallida Eger, 1978 [PER]*

*Loxa peruviensis* Eger, 1978 [PER]

*Loxa planiceps* Horváth, 1925 [DRE]

*Loxa virescens* Amyot & Serville, 1843 [MEX, HON, NIC, PAN, VEZ, SUR, FGU, BRA, ARG, CR, COL]

*Loxa viridis* (Palisot de Beauvois, 1811) [USA, MEX, HON, NIC, DRE, PAN, VEZ, FGU, BRA, ECU, ARG, URU, CR, HAI, CUB, JAM, COL]

#### ***Mayrinia*** Horváth, 1925

Type species: *Loxa curvidens* Mayr, 1864, by original designation.

*Mayrinia brevispina Grazia-Vieira, 1973 [PER, BOL, BRA]*

*Mayrinia curvidens* (Mayr, 1864) [BRA, BOL, PAR, ARG]

*Mayrinia rectidens* (Mayr,1868) [BRA, PER]

*Mayrinia variegata* (Distant, 1880) [NIC, CR, COL, VEZ, GUY, BRA, PER, MEX, HON]

#### ***Rhyncholepta*** Bergroth, 1911

Type species: *Rhyncholepta grandicallosa* Bergroth, 1911, by monotypy.

*Rhyncholepta grandicallosa* Bergroth, 1911 [PAN, VEZ, FG, BRA, CR, HON, COL]

*Rhyncholepta henryi* Kment, Eger & Rider, 2018 [FG]

*Rhyncholepta meinanderi* Becker & Grazia, 1971 [VEZ, BOL, BRA]

*Rhyncholepta wheeleri* Kment, Eger & Rider, 2018 [GUY]

## abbreviations

ARG: (Argentina)
BOL: (Bolivia)
BLZ: (Belize)
BRA: (Brazil)
CHI: (Chile)
COL: (Colombia)
CR: (Costa Rica)
CUB: (Cuba)
CUR: (Curaçao)
DRE: (Dominican Republic)
ECU: (Ecuador)
FG: (French Guyana)
GRE: (Grenadines)
GUY: (Guyana)
GTM: (Guatemala)
HAI: (Haiti)
HON: (Honduras)
JAM: (Jamaica)
MEX: (Mexico)
NIC: (Nicaragua)
PAN: (Panama)
PAR: (Paraguay)
PER: (Peru)
SVG: (Saint Vincent and the Grenadines)
PUR: (Puerto Rico)
SLU: (Saint Lucia)
SUR: (Suriname)
URU: (Uruguay)
USA: (United States of America)
TTO: (Trinindad & Tobago)
VEZ: (Venezuela)

## DATA AVAILABILITY

DNA data are available at GenBank (access codes in Table S2).

## ACKNOWLEDGMENTS

We thank FAPESP for the funding (proc. n. 14/00729-3) and for a PhD Fellowship to BCG (proc. n. 14/21104-1). CAPES and CNPq for PhD Fellowships to CG. CNPq for a Research Fellowship to JG (proc. n. 305009/2015-0). Juliete Costa and Marcel Neves for laboratory technical support. Dept. of Biology at UND for funding, and Matthew Flom and Kenneth Drees for technical support to RBS.

## AUTHOR CONTRIBUTIONS

Designed the project: CG, JC and CFS; Collected molecular data: BCG, CG, SK and RS; Collected morphological data: CG, DAR, JG and CFS. Analyzed the data: BCG and CG; Wrote the manuscript draft: BCG; Contributed to the final version of the manuscript: BCG, SK, DAR, RS, JG and CFS.

## CONFLICTS OF INTEREST

The authors declare no conflicts of interest.

## SUPPORTING INFORMATION LEGENDS

**Table S1.** Genes included in study, primer sequences and sources and locus-specific annealing temperatures (AT).

**Table S2.** Species used as terminal taxa along with their taxonomic classification and availability of molecular markers. X indicates missing data and asterisks indicate novel data.

**Table S3.** Morphological characters with respective references (except for new characters).

**Table S4.** Table of coding for the morphological characters.

**Table S5.** Alignment sizes and results of model selection from ModelFinder to each molecular marker and morphology.

**Figure S1.** Bayesian majority consensus tree constructed from the 69 morphological characters. Numbers above branches are posterior probabilities.

**Figure S2.** Bayesian majority consensus tree constructed using five molecular markers in combination: 16S rDNA, 18S rDNA, 28S D1 rDNA, 28S D3-D5 rDNA and COI mitDNA. Numbers above branches are posterior probabilities.

**Figure S3.** Maximum likelihood tree constructed in IQ-TREE v. 1.6.9 using combined morphological (69 characters) and DNA (16S rDNA, 18S rDNA, 28S D1 rDNA, 28S D3-D5 rDNA and COI mitDNA) data. Numbers above branches indicate support levels estimated through ultra-fast bootstraping.

## Notes

### Competing Interest Statement

The authors have declared no competing interest.

### Summary of Updates

The updated version has an addition to the title, a few minor text edits, and the addition of Fig. 4.

